# Modular core network constructed from *Escherichia coli* transcriptome datasets using a hypergraph-based pan-network approach

**DOI:** 10.64898/2026.09.16.752242

**Authors:** Zhenbo Jiang, Ikuo Uchiyama

## Abstract

We developed a hypergraph-based pan-network framework to integrate transcriptomic co-expression networks across diverse conditions. By applying frequent itemset mining to dataset-specific gene clusters, we constructed a hypergraph capturing frequently co-expressed gene sets. Extracting high-frequency hyperedges yielded a robust core modular network. Applied to 106 *Escherichia coli* transcriptomes, our method accurately recapitulates 70% of known operons within its core modules, outperforming conventional graph-based approaches. Furthermore, through inter-modular network visualization and modularity profiling, this approach successfully reveals the dynamic reorganization of bacterial co-expression architectures in response to environmental changes, offering a powerful tool for large-scale omics integration.

## Background

The rapid accumulation of genomics and other omics data within a given species has advanced our understanding of the diversity and complexity of its genomic structures and functions. The “pan-genome”[1,2] concept first captured this diversity at the genomic scale by defining the pan-genome as the union of all genes present across multiple strains of a species, comprising the core genome (the conserved part) and the accessory genome (the remaining part). This concept has since been extended to the transcriptome level, giving rise to the “pan-transcriptome”[3–5], which represents the entire repertoire of transcripts within a single species. Within this framework, a major challenge lies in integrating quantitative data from heterogeneous transcriptome datasets, as batch effects, variations in sequencing platforms, and inconsistent gene annotations can distort cross-dataset comparisons and mask consistent co-expression structures[6–8].

To address this challenge, the pan-network framework[5,9] was introduced. It first creates a co-expression network for each dataset by connecting gene pairs with strongly correlated expression patterns as edges. Then it forms the pan-network and core networks by taking, respectively, the union and the intersection of the edges from the individual co-expression networks (Figure 1A). In this framework, the universality of an edge (*U*) is defined as the number of datasets in which that edge appears[9], serving as a measure of the commonality across the experimental conditions tested. However, in practice, the vast majority of edges exhibit low *U* (often *U* = 1), meaning that a pan-network is dominated by dataset-specific, possibly noisy or transient interactions. Therefore, extracting a stable ‘core network’ by filtering out low-confidence edges is a crucial approach for capturing biologically meaningful cross-condition patterns.

**Figure 1.**
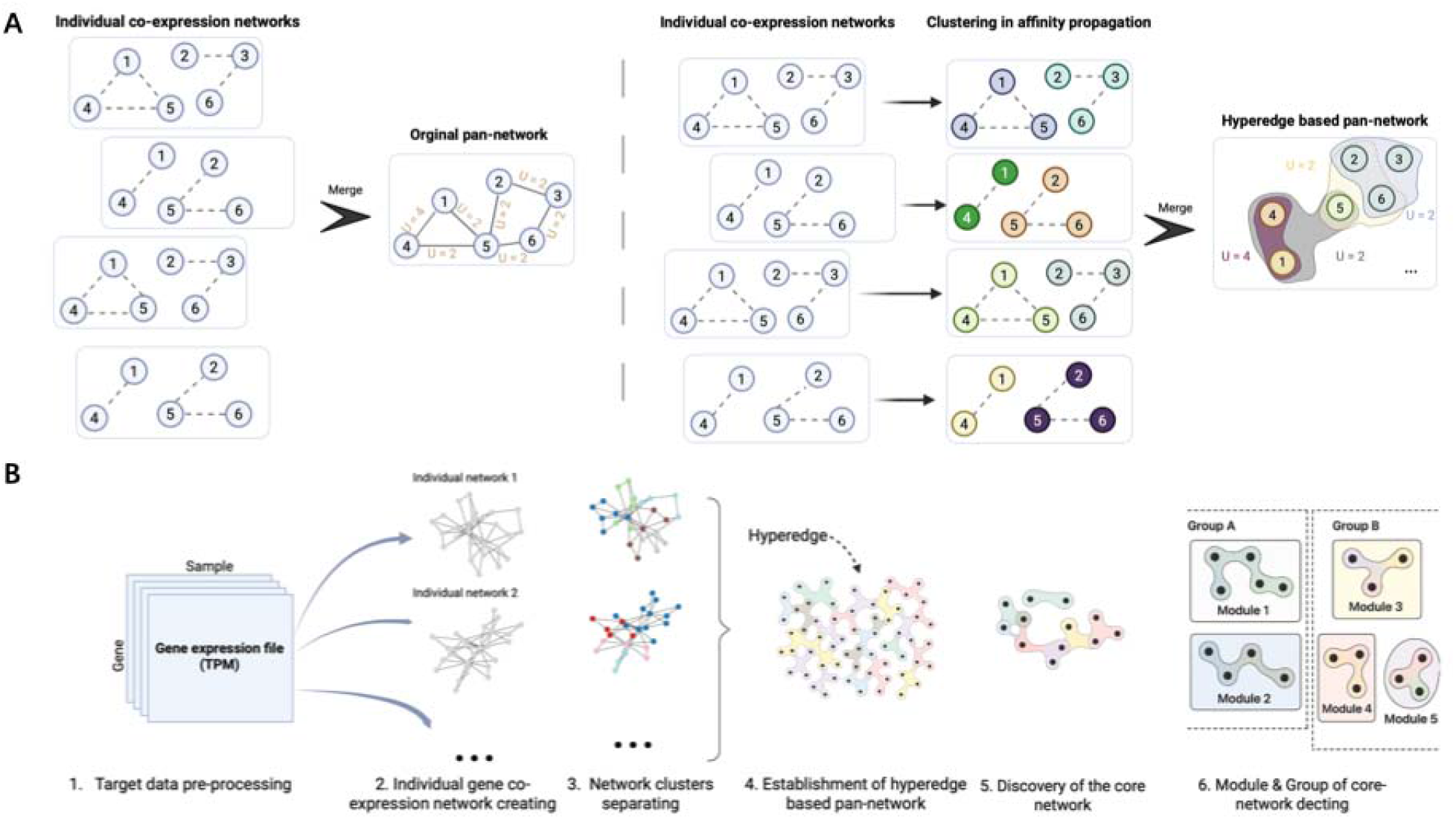
Conceptual framework and construction workflow of the hypergraph-based pan-network. (A) Comparison of traditional edge-based aggregation (left) and the proposed hyperedge-based aggregation strategy (right). In the hyperedge-based approach, clusters of co-expressed genes (nodes) are identified using affinity propagation clustering and then merged across datasets according to recurrence (*U*) to form hyperedges (colored shadows). (B) Schematic overview of the six-step implementation pipeline: (1) transcriptome data preprocessing (TPM normalization); (2–3) construction of individual co-expression networks and cluster separation; (4) assembly of the pan-network by aggregating recurrent hyperedges; (5) identification of the core network based on high *U* values; and (6) decomposition of the core network into biologically coherent modules and groups. Created in https://BioRender.com

Biologically, conserved regulatory patterns rarely manifest as isolated gene pairs; instead, they operate as coordinated, multi-gene units or network modules, such as protein complexes, metabolic pathways, and transcriptional regulons[10,11]. Therefore, one of the ultimate goals of co-expression network analyses is to identify such robust modular structures. In this work, we mainly focus on identifying co-expression modules within the core network of a given species. In this context, despite their simplicity and intuitiveness, conventional graph-based pan-network approaches may fail to accurately capture co-expressed gene sets that recur across datasets. In the example in Figure 1A, two different co-expression patterns ({1,4,5} and {2,3,6}, and {1,4} and {2,5,6}) are observed across the four individual co-expression networks. However, when these networks are merged at the edge level, the gene sets that consistently co-express are broken into separate pairwise links. Consequently, the resulting pan-network loses information about the coordinated expression of multiple genes.

To overcome these limitations, we propose a hypergraph-based pan-network framework (Figure 1B) that elevates co-expression analysis from pairwise edges to gene clusters[12–14]. In our approach, each dataset yields gene clusters, which are then mined across datasets using frequent itemset mining algorithms[15,16] to identify hyperedges—gene sets that are repeatedly co-expressed across multiple datasets. We define the *U* of a hyperedge as the number of datasets in which it appears. The pan-network is then constructed as the union of hyperedges, and the core network consists of those hyperedges that exceed a chosen *U* threshold.

In this study, we focus on bacterial transcriptome data, particularly those of *Escherichia coli* (*E. coli*), to evaluate our extended pan-network method. Bacteria provide an advantageous system for extending the pan-transcriptome framework. The increasing availability of bacterial RNA-seq datasets[17,18], together with their relatively compact genomes and well-organised gene regulatory systems, provides a valuable foundation for transcriptome-wide network analyses. In bacteria, gene expression is commonly coordinated through operons—clusters of adjacent genes transcribed from a common promoter—and regulons, which consist of multiple operons or genes controlled by a single transcription factor[19–21]. *E. coli* is a primary model organism for systems biology and the best-characterized bacterium so far, and its gene regulatory relationships, including operons and regulons, are comprehensively collected into a database[20]. Here, we applied our novel pan-network approach to integrate *E. coli* transcriptome data.

## Results

### Workflow and data overview

Figure 1B provides an overview of our hypergraph-based pan-network construction pipeline. Starting with multiple transcriptome datasets (step 1), we first converted each expression matrix into a co-expression network, thereby capturing gene–gene relationships within individual conditions (step 2). We then decomposed these networks into modular co-expressed gene clusters, which serve as candidate gene sets for cross-dataset comparison (step 3). By integrating these clusters across all networks, we identified recurrent multi-gene combinations—termed hyperedges, and combined them to construct the pan-network as a hypergraph (step 4). Hyperedges supported by more datasets (High *U*) were subsequently used to define the core network (step 5). Finally, the core network was further partitioned into modules, allowing the identification of functionally coherent gene communities (step 6).

We applied this framework to 106 *Escherichia coli* transcriptomic datasets from the Gene Expression Omnibus (GEO) database[22], each comprising at least 15 samples (see Additional file 1, Figure S1A) and originating from multiple *E. coli* strains (the detailed experimental conditions and clade assignments for all datasets are provided in Additional file 2, Table S1). After mapping transcript reads to a unified pan-genome reference and filtering out low-confidence interactions, we constructed an individual co-expression network for each dataset. The resulting networks varied in size, ranging from 1,600 to 7,500 genes and from 1 million to 30 million edges (see Additional file 1, Figures S1B and S1C), depending on the experimental conditions included in the series, such as genetic perturbations and environmental stresses.

To construct the hypergraph-based pan-network, each network was decomposed into clusters using the affinity propagation (AP) clustering algorithm[23], yielding between 60 to 300 clusters across different datasets (see Additional file 1, Figure S1D). Next, we applied a frequent itemset mining algorithm (Eclat-based) to the combined AP clustering results across all datasets to identify hyperedges. Itemset mining identifies groups of commonly co-expressed genes, representing higher-order relationships that are often misidentified by pairwise methods. By setting a minimum cutoff of *U* (or minimum support in data mining terminology), we identified hyperedges in the core network. Finally, for community detection, the hypergraph-based core network was converted into a conventional graph-based core network by replacing each hyperedge with a fully connected subgraph (detailed below).

### The comparison of pan-network methods based on internal and external indices

To evaluate the methods and determine an appropriate cutoff for *U*, we performed benchmark tests. Here, to compare our hypergraph-based pan-network with traditional edge-based approaches, we constructed two benchmark pan-networks. In each dataset, the top 1% and top 5% of gene pairs ranked by pairwise Pearson correlation coefficients (PCCs) were selected to build individual networks, which were subsequently integrated into two traditional pan-networks[9].

We constructed a series of networks with varying *U* and calculated the modularity[24] (Figure 2A) and clustering coefficient[25] (Figure 2B) of each network. Modularity is based on the community structure identified using the Louvain algorithm[26] (a greedy optimization method that maximizes modularity to identify densely connected communities), whereas clustering coefficient quantifies the tendency of nodes to form local clusters without explicitly defining cluster boundaries. In all networks, modularity initially shows a similar increasing trend with *U*, but reaches its maximum at different *U* values (Figure 2A). The clustering coefficient in these networks also exhibits a similar S-shaped trend, reaching its maximum at *U* values similar to those of modularity (Figure 2B). The lower modularity and clustering coefficients observed at low *U* values likely indicate that retaining low *U* edges results in a noise-dominated network, yielding a disorganised structure rather than coordinated transcriptional organization. To confirm this, we also calculated several additional network measures and found that pan-networks with low *U* edges exhibit high density[27] (ratio of actual to possible connections; Additional file 1, Figure S2A), but lack Scale-free topology[28–30], which is indicative of hub-driven network organization observed in typical biological interaction networks (Additional file 1, Figure S2B).

**Figure 2.**
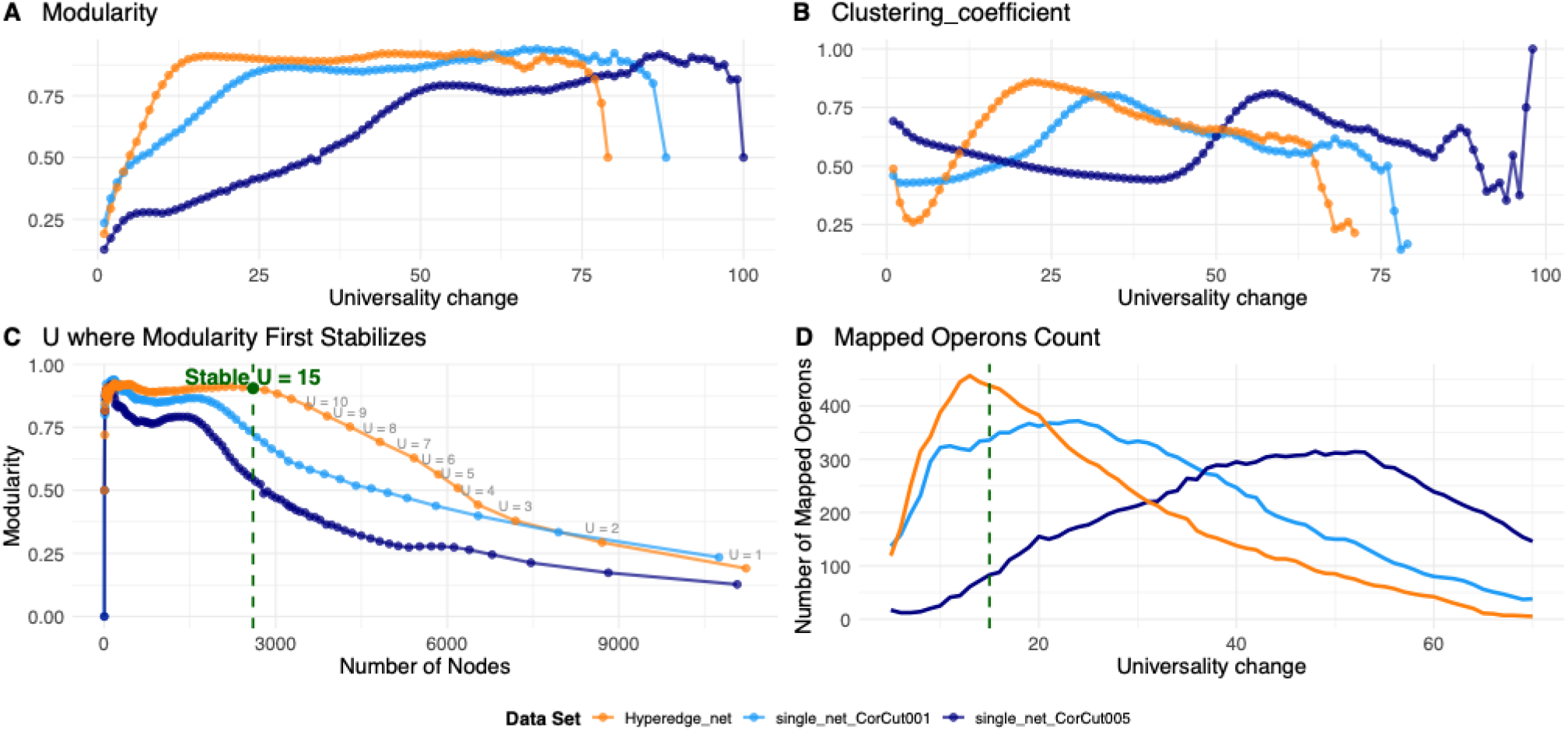
Network characteristics and operon mapping across *U* changes. (A, B) Modularity (A) and clustering coefficient (B) of three network constructions—Hyperedge_net, single_net_CorCut001, and single_net_CorCut005—plotted across increasing levels of U. (C) Modularity as a function of the number of nodes, with *U* values scanned from right to left. (D) The quantity of mapped operons (F-score > 0.5) relative to RegulonDB for the three network constructions. The green dashed line in (C) and (D) marks *U* = 15.

We next evaluated the balance between modularity and network size, measured as the number of non-isolated nodes in the pan-network (Figure 2C). As *U* increases (plotted from right to left), the modularity of the hypergraph-based pan-network rises rapidly, then plateaus at *U* = 15. A further increase in *U* reduces the network size. The two graph-based pan-networks exhibit a similar trend but plateau at lower modularity values and smaller network sizes. Consequently, the hypergraph-based pan-network exhibited a more pronounced modular structure covering a larger set of genes than the graph-based pan-networks. We therefore adopted the hypergraph-based pan-network with a cutoff of *U* = 15 as the optimal core network.

To assess how the detected network modules were congruent with known co-regulated gene sets, we evaluated the overlap between the detected modules and operons defined in RegulonDB[20] using the F-measure metric[31] (Additional file 1, Figure S3A). All networks showed a distinct unimodal trend in the number of mapped operons, with the highest number observed in the hypergraph-based network at approximately *U* = 15 (Figure 2D). In this pattern, the initial increase reflects a reduction in noisy edges, similar to the trend observed in modularity, whereas the subsequent decline likely reflects network fragmentation. This fragmentation is evident in changes in two quality metrics for each operon: while precision (Additional file 1, Figure S3A) remains ≥ 0.9 on average at higher *U* values, recall(Additional file 1, Figure S3B) declines substantially with large variance, indicating that some operons are divided into smaller co-expression modules at higher *U* values.

### Core network communities detection and annotation

We identified 178,103 hyperedges (frequent closed itemsets identified by data mining; see Methods) involving 2,604 genes in the hypergraph-based core network (*U* ≥ 15). The resulting hypergraph was subsequently converted into a standard graph representation of the core network (Figure 3) by replacing each hyperedge with a fully connected subgraph. Then, we performed community detection on the core network using the Louvain algorithm, as implemented in Gephi[32].

**Figure 3.**
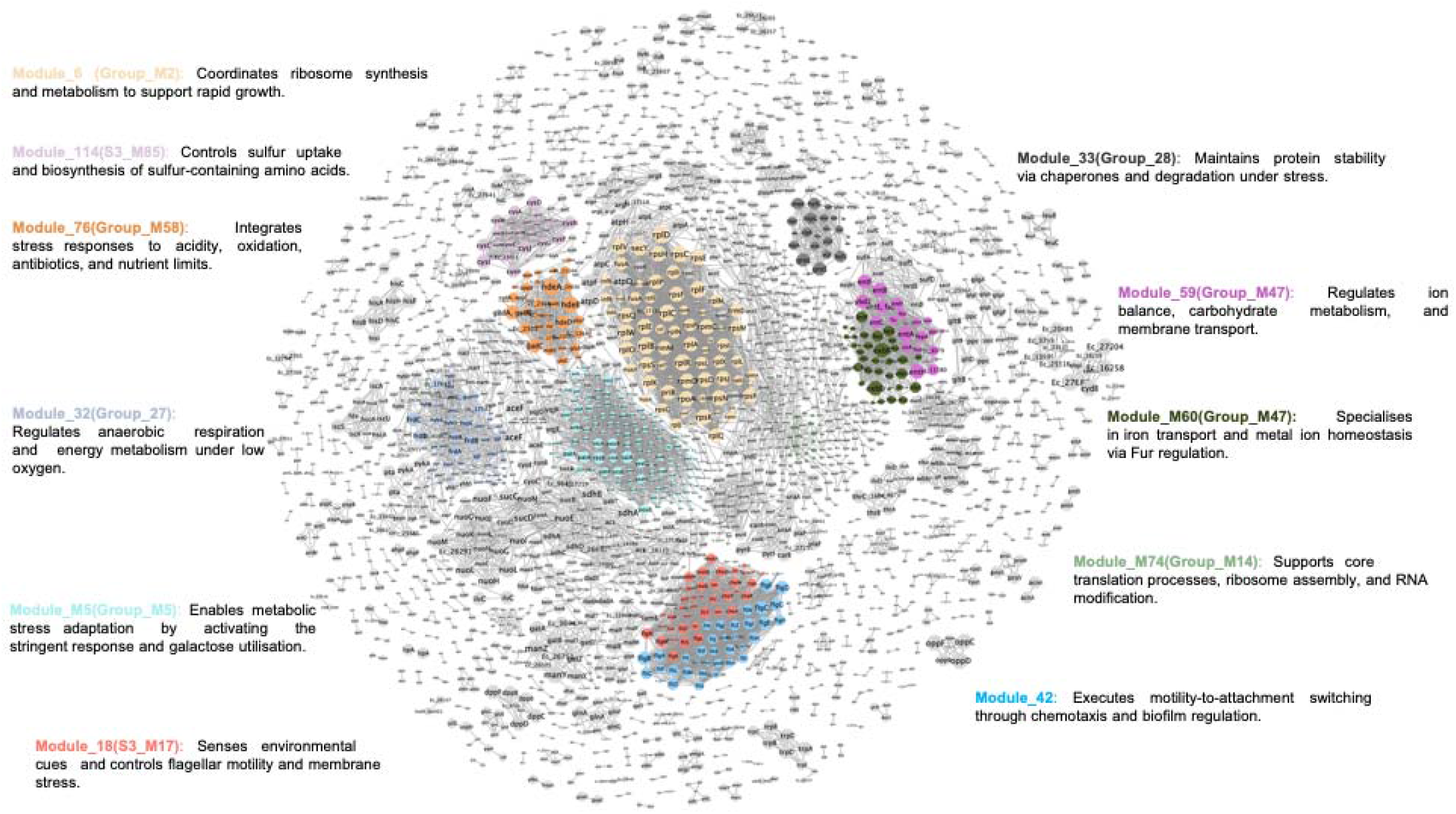
Topological architecture of the core network and spatial distribution of the 11 largest modules. The global structure is visualised using the Compound Spring Embedder (CoSE) layout from Cytoscape. Nodes represent genes, and edges represent interactions. The top 11 largest modules are highlighted in distinct colours to visualise their clustering. Brief functional annotations are provided around the network to summarise the key biological roles of these major modules, as derived from HTable S2. Grey nodes indicate genes belonging to smaller modules.

The Louvain algorithm identifies a community structure with multiple levels, and here we focus only on two levels: the lowest-level communities as ‘Modules’ and the highest-level communities as ‘Groups,’ which are composed of multiple modules (note that not all modules are incorporated into groups). By benchmarking these communities against RegulonDB functional terms (Additional file 1, Figure S4), we found that the module size distribution closely mirrors that of known operons, suggesting that these modules represent fundamental, tightly knit transcriptional units. In contrast, the Groups display a size distribution comparable to that of regulons, suggesting that they are associated with broader biological processes or intricate regulatory networks.

We characterised all modules containing ≥ 5 genes using functional enrichment analysis with RegulonDB as well as functional classification in MBGD (see Additional file 2, Table S2 and S3). Here, we focus on the 11 largest modules highlighted in Figure 3, along with their functional annotations. The two largest modules are Module 5 and Module 6, which are identified as a regulatory unit tightly linked to the ‘ppGpp’ regulon and a ribosomal complex, respectively (see Additional file 1, Figure S5 for the relationships between the largest modules and their associated regulons and functional categories represented as a bipartite graph). Modules show contrasting *U* profiles that determine their topological complexity (Additional file 1, Figure S6A). Notably, within-module hyperedge counts do not scale linearly with module size (Additional file 1, Figure S6B). For example, Module 6 (75 genes, high U) has >100,000 hyperedges, while the larger Module 5 (138 genes, low U) has only ∼1,000. This disparity results from structural differences between closed and maximal itemsets (Additional file 1, Figure S6C). Specifically, closed itemsets include all observed hyperedges, while maximal itemsets are the non-redundant ones selected from these hyperedges. Modules with lower *U* values (e.g., Modules 5, 32, 60, 74) show minimal redundancy, align with the diagonal, and are mainly composed of pairwise edges. Conversely, high *U* modules (e.g., Module 6) exhibit extensive combinatorial redundancy via many hyperedges (>2 genes), indicating that high *U* links to multigene co-expression complexes, whereas low *U* relates to a looser pairwise interaction topology.

Among the largest modules, two neighbouring modules are related to cell motility and exhibit different *U* profiles: Module 42 shows high *U*, while Module 18 shows a bimodal distribution, suggesting a hybrid composition of essential and accessory components. Module 42, comprising about 40 genes, contains over 18,000 hyperedges. This suggests that distinct gene combinations are co-expressed under varying conditions, reflecting the complexity of underlying interactions. Another pair of strongly related modules is Module 59 and 60: Module 60 on siderophore transport has a wide *U* distribution, while Module 59 on ferrous iron transport has a narrower distribution, forming ‘Iron Homeostasis’ units.

### Relationships Among Sub-communities Within the Core Network

To evaluate the biological relevance of the identified modules, we mapped them to regulatory units defined in RegulonDB, including operons and regulons, and additional units defined based on their sets of regulating transcription factors (TFs): “regulon combination strict” (RC_strict), defined as a set of genes regulated by exactly the same set of multiple TFs, and “regulon combination extended” (RC_extended), defined as a set of genes regulated by multiple common TFs (see Methods). The proportion of each mapping state is summarised in Figure

4. Most identified modules (71.3%) were mapped to operons, consistent with their size distribution (Additional file 1, Figure S4). Many (n=285) had perfect agreement with known operons (‘Perfect Match’), while others (n=86) were subsets (‘Module is Subset’), indicating alternative start or stop sites. Fewer modules matched RC categories (RC_extended: 4.7%; RC_strict: 10.5%) or Regulons (5.7%). About 7.8% remained unmapped, likely representing non-*K-12* orthologs outside RegulonDB or unannotated *K-12* units.

Building upon these modules as local regulatory units, we sought to visualise how they assemble into the global connectivity of the *E. coli* pan-network modules. To this end, we constructed a module-level network (Figure 5A), where nodes represent modules and edges represent hyperedges connecting them. These connections are further categorised into intra-group (brown) and inter-group (green) hyperedges. Within this network, the 11 largest modules (excluding Modules 59 and 60) are distributed across different groups, and some of them (e.g., Modules 6, 5, and 32) serve as highly connected structural hubs.

**Figure 4.**
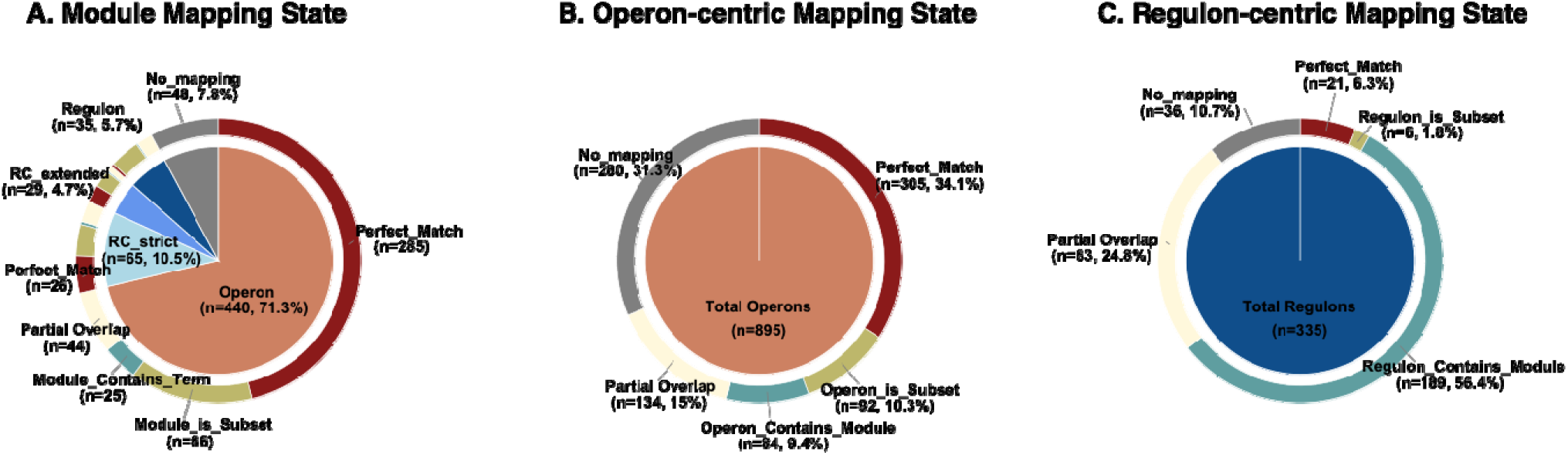
Bidirectional mapping evaluation between pan-network modules and RegulonDB annotations. The triple-panel double-ring pie charts illustrate the functional mappings from the perspectives of network modules (A), known operons (B), and known regulons (C). Inner rings display the total entities evaluated, and outer rings show the distribution of mapping states. Notably, the assigned state for each entity represents its single optimal match found against the respective counterpart background. Mapping categories are defined by F-score and inclusion metrics: Perfect Match (F-score ≥ 0.8); Subset relations (the entity is strictly contained within the target, Precision/Recall ≥ 0.8); Containment relations (the entity broadly encapsulates the target, Precision/Recall ≥ 0.8); Partial Overlap (overlap count ≥ 2 genes); and No mapping. This bidirectional approach highlights both the biological composition of the identified modules and the recovery rate of established regulatory structures.

**Figure 5.**
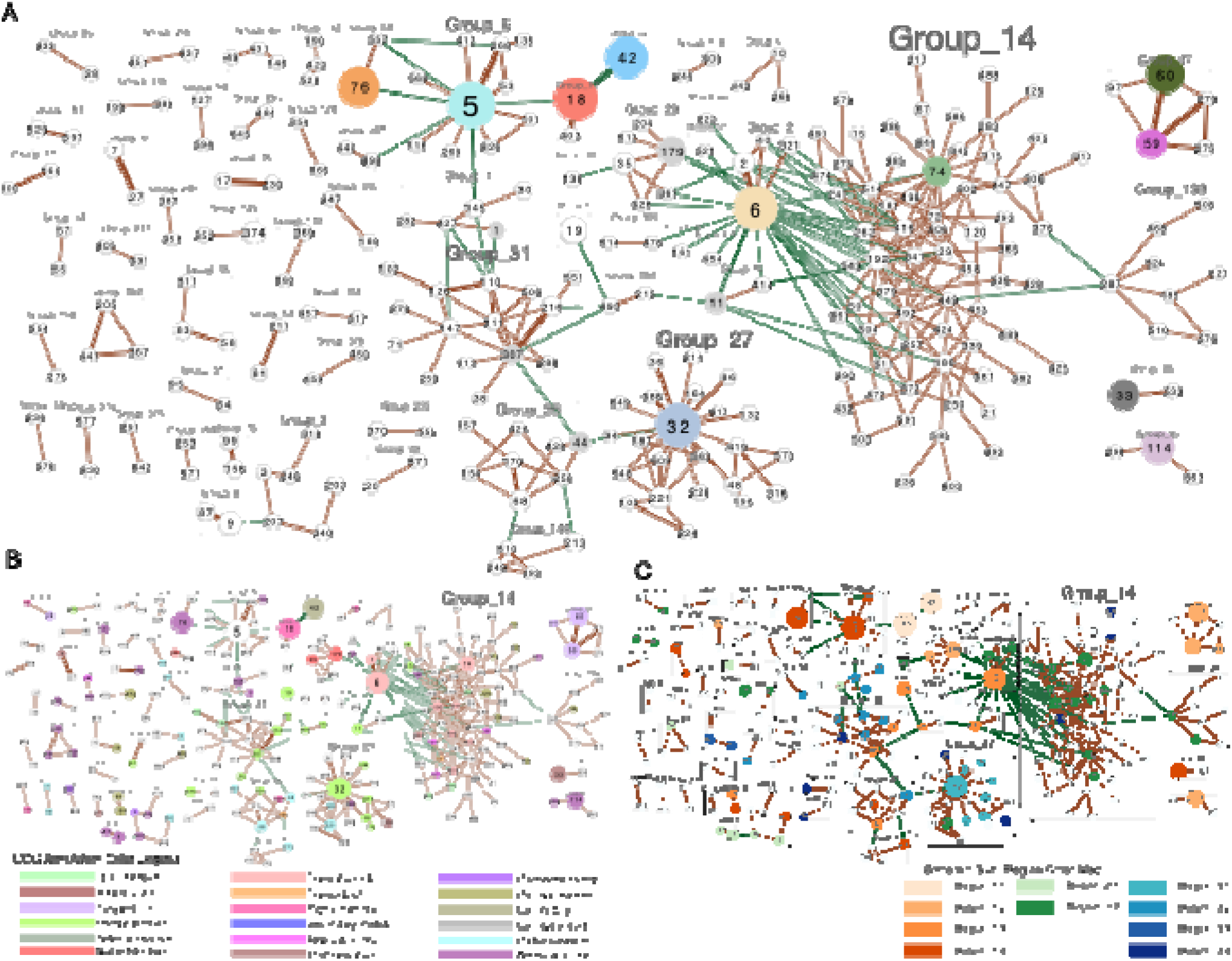
Inter-modular connectivity within the core-network. (A–C)Network visualization of modules connected by shared hyperedges. Nodes represent modules (labelled by ID, sized by gene count), and shaded backgrounds delineate distinct Groups. Edge width reflects the number of shared hyperedges (green: inter-group; brown: intra-group). (A) Topological overview. The top 11 largest modules are distinctly colored; five selected modules of interest are highlighted in grey. (B) Functional annotation. Node colours indicate the primary COG category, or the dominant COG class covering >80% of the module genes if a primary annotation is absent; detailed colour mapping is provided in Additional file 2, Table S4. (C) Regional distribution. Node colours denote the Subregion assignment (Regions 1-3) for each module.

Based on the primary COG functional category assigned to each module and the underlying network structure, we identified several hub-centered domains with connections spanning multiple groups (Figure 5B). One domain is centered on the translational hub with the highest degree, Module 6 (ribosomal proteins, *rpl*/*rps* operons), in Group 2. Although it exhibits many connections across multiple groups, most are within Group 14, the largest, with modules such as Modules 74, 192, and 341 dedicated to tRNA aminoacylation and rRNA modification, thereby supporting translation, while modules 120, 488, and 514 focus on cell wall synthesis and morphology.

Another domain comprises groups primarily associated with energy metabolism, including Groups 27 (Anaerobic respiration), 31 (TCA cycle), 36 (Carbohydrate transport and metabolism), and 158 (Energy production and conversion). Unlike the previous domain, where inter-group connections are primarily concentrated on the central hub module, these groups are interconnected by cross-group links mediated by several specialized nodes, including Modules 1, 11, 179, 307, and 44. For example, Module 307 mediates aerobic carbon processing in Group 31, linking the TCA cycle to Group 36’s carbohydrate transport via Module 44 (responsible for glycolysis and PTS-mediated sugar uptake), and to Group 27 (anaerobic respiration) via Module 34 (associated with nitrogen metabolism and nitrate respiration).

The third domain is built around Module 5, the network’s largest entity, characterized by the ppGpp regulon. This module is connected via cross-group links to Module 18 (chemotaxis genes), Module 42 (flagellar operons), and Module 76 (acid-resistance operons), suggesting that this domain coordinates stress response and motility.

### Bipartite relationship between datasets and core network modules: Invariant backbone vs. Adaptive plasticity

To evaluate the activation of core network modules across 106 transcriptomic datasets, we quantified the normalised Modularity[24] (*Qc*, see Methods) for modules with size ≥ 5, a metric to evaluate modularity of individual modules by decoupling the intrinsic scaling effect of module size, and constructed the module activation matrix (Figure 6). Bidirectional clustering of the matrix revealed three distinct module regions (Regions 1–3, comprising 10 subregions) and three distinct dataset clades (Clades 1–3) (Figure 6A). This bipartite mapping reveals a clear gradient in activation strength across both dimensions: R1 > R2 > R3 for module regions and C1 > C2 > C3 for dataset clades. Notably, Region 1, the region with the highest modularity, contains 9 of the 11 largest modules listed above, whereas the remaining regions harbour only one each.

**Figure 6.**
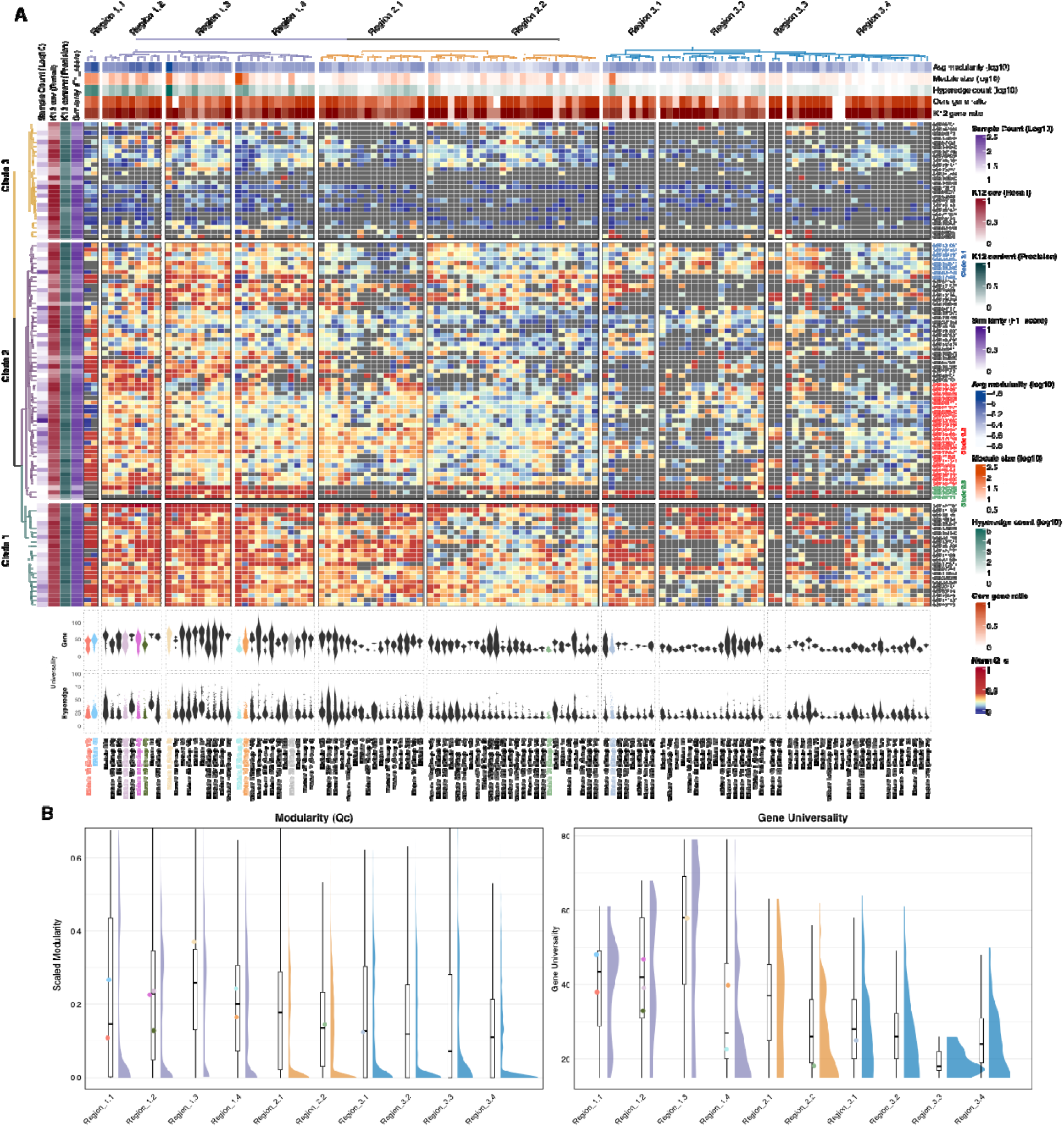
The modularity profile of the *Escherichia coli* core-network. (A) Bi-directional clustered heatmap of normalized module efficiency (Modularity Qc). Rows represent 106 transcriptomic datasets clustered into three major Clades (Clades 1-3), reflecting distinct module states across the datasets. Columns represent core network modules clustered into major Regions (Regions 1-3) and 10 subregions based on their activation patterns. Left annotations indicate dataset-specific validation metrics: Log10 sample count, *E. coli K-12* genome coverage (Recall), genome content (Precision), and network similarity (F1-score). Top annotations display intrinsic module properties: Average modularity (log10), module size (log10), hyperedge count (log10), and core gene ratio. (B) Raincloud plots depicting the statistical distribution of Scaled Modularity (left) and Gene *U* (right) across the 10 subregions. Each raincloud features a half-violin plot (distribution density), a central narrow boxplot (quartiles), and jittered scatter points (Colors correspond to the top 11 modules in Figure 3), representing the median values of individual modules within that subregion. Half-violin colors correspond to the parent Region mapping in Figure 6 (Region 1: purple, Region 2: orange, Region 3: blue).

To deconstruct their organisation, we divided these regions into subregions (Figure 6A). These subregions exhibit distinct distributions of modularity and gene *U* (Figure 6B), with gene *U* defined as the highest *U* score among the hyperedges each gene participates in. Mapping these regions onto the global modular network reveals that they tend to form clusters (Figure 5C), suggesting that they correspond to clusters of co-activated modules. Based on this mapping (Figure 5C), along with functional enrichment analysis with the KEGG pathway categories (see Additional file, Table S3) and the intersection of shared regulators (Additional file 1, Figure S7), the specific characteristics of each region in Figure 6A are detailed below: Region 1 (n = 33) includes four subregions with high *U*. R1.1 consists of two modules (18 and 42) associated with cell motility and chemotaxis. R1.2 comprises clusters of Groups 47 (iron homeostasis, including Modules 59 and 60) and 29 (nucleotide metabolism). R1.3 covers the core translational Module 6 (translational hub) and several modules involved in central energy metabolism, including Module 11 (ATP synthesis) and 307 (TCA cycle). R1.4 involves Modules 5 and 76, stress-adaptation networks regulated by ppGpp and GadX/W. Region 2 (n=42) is divided into two subregions with intermediate *U*. R2.1 comprises modules in Group 3 (polysaccharide and amino acid metabolism) and Group 8 (methionine biosynthesis), with several other modules scattered across independent groups. Meanwhile, R2.2 harbours numerous modules connected to Module 6, the translation hub, which mainly belong to Group 14 and 2. Region 3 (n=48) exhibits four subregions with the lowest *U*. R3.1 is predominantly composed of modules in Group 27 (anaerobic respiration), including Module 32. R3.2 spans Groups 1, 31, and 36, encompassing modules involved in carbohydrate and secondary carbon metabolism under CRP control. R3.3 consists of two modules in Group 46 that encode operons involved in cell wall and capsule biosynthesis. In contrast, R3.4 is scattered across distinct network regions, although some of its modules maintain connectivity with other subregions within Region 3.

The module activation matrix (Figure 6) also shows that modularity values for the same module vary across datasets; based on these values, the datasets are classified into three clades (Clades 1-3) (Additional file 2, Table S1). Clade 1 exhibits the highest modularity, whereas Clade 2 displays intermediate levels. In contrast, most modules in Clade 3 are silenced or exhibit minimal modularity. Based on these observations, the specific characteristics of each clade are detailed below: Across individual transcriptomes in Clade 1, core network modules robustly preserve their distinct boundaries, as evidenced by uniformly high modularity (Figure 6A). The datasets within this clade are derived from *E. coli K-12* and related strains in the exponential growth phase, under various experimental conditions. These conditions include active metabolic optimization via adaptive laboratory evolution (ALE) [e.g., GSE114358, GSE97944] and targeted regulatory evaluations under mild perturbations [e.g., GSE143855, GSE225096]. Clade 2 (n = 57) exhibits intermediate modularity yet comprises highly heterogeneous datasets, including those exposed to diverse environmental transitions and sublethal stresses. Within this broad, heterogeneous clade, we focus on three subclades that exhibit distinct topological and biological features in the network matrices (Figure 6A). Subclade 2.1, indicated by blue labels in Figure 6A, is predominantly composed of transcriptomes derived from cells entering the stationary phase. In addition, it captures related extreme survival transitions, such as cold shock (e.g., GSE152619) or broad-spectrum antibiotic exposure (e.g., GSE220559). Specifically, within Region 3, which contains adaptive stress modules, these datasets exhibit elevated internal density coupled with notably low modularity. Subclade 2.2, indicated by red labels, constitutes a large cluster encompassing datasets from cells subjected to sublethal antibiotic exposure (e.g., GSE184365), synthetic biology payloads (e.g., GSE168336), or structural growth arrest (e.g., GSE144604). This group is characterised by high internal density yet conspicuously low modularity within Region 2, which contains macro-assembly and structural biosynthesis networks. Finally, Subclade 2.3, indicated by green labels, captures a highly specific state of profound epigenetic or physical crisis, such as the genetic deletion of the global repressor H-NS (e.g., GSE40313) or acute antimicrobial peptide damage (e.g., GSE160082). In stark contrast to other subclades, this group features the localized, hyper-intense activation of Subregions R1.3 (including the core translational hub) and R1.4 (including the ppGpp-driven stress adaptation network), while the peripheral structural and adaptive regions remain exceptionally sparse. Clade 3 (n = 26) shows a notably fragmented activation pattern, characterised by the lowest overall modularity and a widespread breakdown of network structure across all regions. Genomically, the datasets in this group contain a higher proportion of non-K12 pan-genomic features than those in Clades 1 and 2. These transcriptomes mainly reflect bacteria experiencing severe survival challenges, including acute genetic disturbances [e.g., GSE102381, GSE168963], lethal phage infections [e.g., GSE161794, GSE96573], and intense environmental stresses [e.g., GSE208658, GSE103421]. Under these extreme physiological stresses, the core metabolic system and overall modular structure break down severely.

To understand the implications of low modularity, we subsequently calculated the network density (the ratio of actual to possible connections) within each module (Additional file 1, Figure S8). Crucially, Clade 2 generally maintained internal module densities comparable to those in Clade 1, whereas Clade 3 exhibited lower densities. This suggests that the lower modularity in Clade 2 is not due to the disruption of within-module co-expression relationships but rather to an increase in cross-module co-expression interactions. To confirm this, we categorized edges into four categories: inner-module (IM), cross-module/inner-group (IG), cross-group (CG), and accessory (Acc). We then compared the proportions of edges in each category across datasets within Clade 1 and Clade 2 of the same module (Additional file 1, Figure S9A). The results indicate that the primary difference lies in a significantly higher abundance of CG and Acc edges in Clade 2 than in Clade 1, whereas the differences in IM and IG edges are confined to a limited subset of modules (with a few exceptions in IG edges; see Discussion). This suggests that substantial network rewiring occurs in Clade 2 outside the defined Groups.

## Discussion

Pan-network analysis effectively integrates diverse transcriptomic datasets, yet constructing biologically accurate networks remains a computational challenge. Previous pairwise approaches[9] are highly susceptible to transitive correlations across datasets, obscuring genuine co-expression relationships (Figure 1A). Our hypergraph-based formalism overcomes this by capturing simultaneous, multi-way co-expression relationships. As demonstrated in Figure 2 and Additional file 1, Figure S4, this approach more effectively filters stochastic noise, yielding a superior modular structure at lower *U* than conventional graph-based methods, thereby successfully reconstructing core modules with greater coverage of known operons. The roughly 70% reciprocal overlap between our 618 identified core modules and known RegulonDB operons (Figure 4) confirms that our analysis accurately captures operon structures as the fundamental building blocks of the core network.

While core modules encode discrete basal functions, bacterial survival in fluctuating environments requires these units to coordinate under stress. In our approach, these higher-order interactions primarily manifest as a hierarchical modular structure (Group) and are further visualized as an inter-module network based on the underlying hyperedges (Figure 5). In addition, the modularity profile (Figure 6) delineates the activation states of individual modules across diverse experimental conditions, thereby revealing higher-order cooperative dynamics among them.

Recently, matrix decomposition methods, such as independent component analysis, have been applied to integrate transcriptomic data across diverse conditions[9,33]. Although powerful, these approaches require careful experimental design under homogeneous conditions or rigorous correction of batch effects. In contrast, our pan-network approach effectively integrates heterogeneous transcriptomic datasets without requiring batch-effect correction. Indeed, the successful incorporation of all available datasets—including those derived from severe or even highly artificial conditions (represented in Clade 3)—robustly demonstrates the technical resilience and versatility of our method.

Within the modularity profile (Figure 6), *E. coli* core network modules are classified into three distinct regions: Region 1 features highly independent activation units that are commonly activated across a broad range of datasets (high *U* and modularity; dense intra-but sparse inter-module connections); Region 2 contains modules prone to coordinate co-activation (lower *U*; dense connections within and across modules); and Region 3 consists of modules that are sporadically activated in a condition-specific manner. The modularity profile also captures dynamic hyperedge crosstalk across modules in these regions, enabling bacteria to adapt to various experimental conditions. In this work, the transcriptomic changes in the available *E. coli* datasets are categorized into three distinct clades: Clade 1 represents baseline physiological maintenance under standard growth[17,34]; Clade 2 exhibits heterogeneous adaptation to environmental transitions and sublethal stresses[35]; and Clade 3 represents systemic network breakdown under severe survival challenges, such as lethal phage infections or acute genetic disturbances[36].

Our hyperedge framework captures the physiological rewiring driving this adaptation. Specifically, while Region 2 modules exhibit comparable inner-module densities between Clade 1 and 2 (Additional file 1, Figure S8), their overall modularity drops significantly in Clade 2 (Figure 6). Under favorable Clade 1 conditions, these functional modules act with high independence, requiring few inter-module connections. In contrast, the sublethal stress conditions of Clade 2 trigger widespread systemic defense responses. For example, Region 2 modules in Group 14 are linked to defensive processes (Additional file 2, Table S5), including suppression of macromolecular synthesis—such as DNA replication and translation—(e.g., the replication helicase *dnaB*[37] and stringent response regulator *spoT*[38] in Module 74, and the ribosomal silencing factor *rsfS*[39] in Module 514) and reinforcement of the cell envelope. Crucially, our network identifies these responses at the level of functional complexes and operons rather than at the level of isolated genes. This is shown by the BAM complex and its associated genes (*bamA, bamB, rseP* [Module 186])[40], the enterobacterial common antigen (ECA) biosynthesis gene cluster (*wec/rff* operon, including *wecB* and *wecC* [Module 120])[41], and the peptidoglycan synthesis operon *mrdAB* (*mrdA, mrdB* [Module 514])[42]. Coordinated activation of these modules accounts for the observed increase in intra-group connections among them, as well as the rise in cross-group and accessory connections (Additional file 1, Figure S9B). Therefore, the observed decrease in modularity in our data biologically reflects a rewiring of the transcriptional network, facilitating a reallocation of transcriptional resources, a known mechanism that boosts resilience and survival under sublethal stress[30,43–46]. Ultimately, when extreme perturbations exceed these buffering limits (Clade 3), this coordinated system collapses into widespread silencing and systemic fragmentation[35,45].

The developed modular core network can serve as a reference framework for comparative transcriptomic analysis. A straightforward application is to map new *E. coli* transcriptomic datasets, obtained under different conditions or from different strains, onto the core network modules using representations such as the inter-module network (Figure 5) or the modularity profile (Figure 6), thereby facilitating their characterization. A more challenging application is cross-species transcriptome comparison. Since the core network encompasses a broad range of transcriptomic states across diverse datasets, it provides a useful framework for comprehensive cross-species transcriptome comparison by enabling the identification of conserved and species-specific network features from an evolutionary perspective[47].

In this work, we utilized all available *E. coli* transcriptome datasets to construct the pan-network. Although we successfully delineated core modules even from such highly heterogeneous datasets, this heterogeneity limits the interpretability of the dataset-modularity profile with respect to dataset-specific network reorganization. This limitation can be overcome by integrating our framework with a systematically designed experimental series. Recent advances in single-cell RNA sequencing (scRNA-seq) technologies have enabled the development of several tools to extract individual networks, such as cell-type-specific networks, from scRNA-seq datasets[48,49]. Integrating these individual networks is another promising application of our framework.

## Conclusions

Our hypergraph-based pan-network framework is a computational methodology that explicitly models higher-order, multi-gene co-expression relationships across heterogeneous transcriptomic data to achieve large-scale omics integration without requiring strict batch-effect correction, core network reconstruction and module identification. The framework leverages frequent itemset mining on dataset-specific gene clusters to capture recurrent multi-gene combinations, and extracts high-frequency hyperedges to construct a robust core network that accurately recapitulates fundamental known operon structures. It also supports interpretable analysis of dynamic network rewiring through modularity profiling and enables downstream tasks, such as core regulatory unit identification and stress adaptation analysis. Moreover, our approach exhibits robust technical resilience, delivering precise insights into both invariant metabolic backbones and highly plastic adaptive modules across diverse experimental conditions, with broad potential for generalization to cross-species evolutionary comparisons and single-cell RNA sequencing (scRNA-seq) analyses.

## Methods

### Data collection and pan-genome reference construction

Expression datasets with at least 15 samples were obtained from NCBI GEO, yielding a compendium of 106 E. coli datasets spanning diverse strains (e.g., *K-12, B*, and clinical isolates). A non-redundant pan-genome reference was constructed using the Microbial Genome Database (MBGD)[50]. Specifically, a representative nucleotide sequence for each *E. coli* orthologous cluster (Taxonomy ID: 562), originally defined using DomClust[51], was retrieved from the MBGD database to serve as a universal reference for cross-strain read mapping. This comprehensive pan-genome reference comprises 27,727 orthologous groups, establishing a standardized set for transcriptome analysis that captures both the core and accessory components of the *E. coli* pan-genome. The details of these datasets are listed in Additional file 2, Table S1.

### High-throughput expression profiling and gene filtering

Transcript abundance was quantified using Salmon (v1.0.3)[52], utilizing the constructed pan-genome as the reference index. Expression levels were calculated and reported as transcripts per million (TPM). Genes with a maximum expression of TPM ≤ 5 across all samples were removed. PCCs were computed for all gene pairs, and genes with no significant co-expression with any other gene (FDR-adjusted *p*-value < 0.05, computed via the Benjamini-Hochberg procedure)[53] were excluded prior to network construction.

### Co-expression network construction and clustering via Affinity Propagation

An independent gene co-expression network was constructed for each of the 106 datasets. Each network was generated using absolute PCCs for all gene pairs, with non-significant correlations (adjusted p-value > 0.05) set to zero. Gene clustering was performed using the Affinity Propagation algorithm. Clusters and exemplars were generated through iterative message passing, utilizing a preference parameter (quantile) of 0.5. Unlike methods such as k-means or hierarchical clustering, AP automatically determines the number of clusters by selecting representative “exemplar” genes based on pairwise similarities. This allows the detection of clusters with varying sizes and connectivities, identifying both dense and sparse co-expression modules within the same dataset. This flexibility is crucial for robustly handling highly diverse transcriptome datasets.

### Pan-network hypergraph construction and pairwise reduction

To construct the pan-network, AP-derived gene clusters from each dataset were treated as distinct transactions. Frequent closed itemsets were mined using the Eclat algorithm (arules R package, v1.7-9)[15,54] to define pan-network hyperedges. Each hyperedge was characterized by its *U*, defined as the number of datasets in which its constituent genes co-occurred within the same AP cluster (i.e., support). To facilitate subsequent community detection and topological analysis, the hypergraph was transformed into a pairwise co-expression network via clique expansion (Additional file 1, Figure S10). Specifically, each hyperedge was decomposed into a fully connected subgraph, with every edge inheriting the *U* value of the original hyperedge. For gene pairs present in multiple overlapping hyperedges, the corresponding edge was assigned the maximum *U* value across all hyperedges, thereby preserving the strongest co-expression signal.

### Core gene identification and community detection

The *E. coli* core genome identified using the CoreAligner program[55] was obtained from the MBGD database. Conservation profiles for all *E. coli* (Taxonomy ID: 562) orthologous clusters were retrieved, with core genes explicitly defined based on the CoreAligner algorithm output[50]. Community detection for both the pan-network and the core network was performed using the Louvain algorithm[26]. To systematically capture community structures, the first output level was defined as fine-grained “Modules,” and the final aggregated level as broader “Groups.” Edge weights were strictly determined by their assigned *U*. Topological properties of the pan-network, including the average clustering coefficient, were calculated using the igraph R package (v1.5.1), whereas modularity analysis of the core network was implemented via Gephi (v0.10.1)[32].

### Functional annotation and mathematical formulation of regulatory combinations

Transcriptional regulatory data were retrieved from RegulonDB (v12.0)[20] and classified into four categories: (1) Operon; (2) Regulon; (3) RC_strict (genes co-regulated exclusively by a fixed combination of transcription factors [TFs]); and (4) RC_extended (genes regulated by subsets of ≥ 2 TFs derived from larger combinations of ≥ 3 TFs in RC_strict). Functional annotations, including COG, KEGG, GO, and MBGD terms, were obtained from the MBGD database.

Let *G* be the set of all genes, and *T* be the set of all transcription factors (TFs). Let *R* ={(t, g)∣t ∈ *T*, g ∈ *G*} denote the set of all directed regulatory interactions. For each gene *g* ∈ *G*, the set of TFs regulating *g* is defined as:

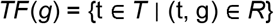

Let *S* = {TF(g)∣g ∈ G} be the collection of all unique TF combinations observed in the regulatory network. For each *S* ∈*S*, the associated strictly co-regulated gene set is defined as:

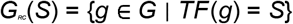

For any combination where *G*_*RC*_(*S*) ≠ ∅, the gene set *G*_*RC*_(*S*) constitutes an RC_strict module. To capture higher-order, nested regulatory architectures, RC_extended sets were derived from RC_strict combinations involving three or more TFs. For a given TF subset *S*′′⊆ *S* (subject to the constraint ∣*S′*′∣≥2), the expanded co-regulated gene set is defined as:

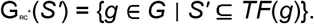

If G_RC_ ^+^+(*S′*) ≠ ∅, it constitutes an RC_extended module, representing genes jointly regulated by at least the TFs in S′, irrespective of additional regulators.

Functional enrichment analysis and module evaluation metrics

Functional annotations of gene modules were mapped and evaluated to determine biological relevance. The concordance between predicted gene modules (M) and actual functional terms (T) was evaluated using Precision, Recall, and F-score[31], formulated as follows:

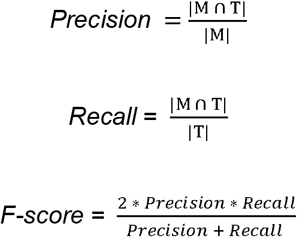

### Network visualization

The core gene network was visualised using Gephi (v0.10.1). Network layout was computed using the Fruchterman-Reingold algorithm[56], with pairwise edges derived from hyperedge reduction.

### Quantification of dataset-specific modularity performance

Condition-specific co-expression networks were constructed by retaining the top 5% significant interactions (FDR < 0.05). Modules with < 5 genes globally or < 3 genes mapped to nodes locally were excluded. The structural performance of each module was quantified using a weighted version of Newman’s Modularity (Qc)[24]. To rigorously eliminate the intrinsic resolution limit and size-induced scoring biases inherent to Qc[57,58], an empirical power-law decoupling was applied. The size-independent modularity was calculated as Qc/(Size^1.4439), where 1.4439 is the empirically derived scaling factor (Additional file 1, Figure S11). Inactive modules (Qc < 0) were designated as NA. Finally, the size-decoupled matrix was globally Min-Max normalized and clustered (Spearman correlation, Ward.D2 method) to systematically partition modules into Regions 1–3 and datasets into Clades 1–3.

### Statistical Analyses

Functional enrichment for modules, groups, and regions was determined using Fisher’s exact test via the clusterProfiler package[59], evaluating terms across seven functional and regulatory categories: Regulon, Operon, KEGG, COG, TIGR, Kmodule, and MBGD. To ensure high confidence, terms were filtered with a minimum module size (≥5 genes) and FDR < 0.05. *P*-values were adjusted for multiple hypothesis testing using the Benjamini-Hochberg (BH) procedure.

### Web Sites/Data Base Referencing

- NCBI GEO: https://www.ncbi.nlm.nih.gov/geo/
- MBGD: https://mbgd.genome.ad.jp/
- RegulonDB (v12.0): https://regulondb.ccg.unam.mx/
- Salmon (v1.0.3): https://combine-lab.github.io/salmon/
- Gephi (v0.10.1): https://gephi.org/
- R packages utilized include *igraph* (v1.5.1) and *arules* (v1.7-9).

## Additional files

**File name:** Additional file 1

**File format:** .pdf **Title of data:** Supplementary Figures for Modular Core Network Constructed from *E. coli* Transcriptome Datasets Using a Hypergraph-based Pan-network Approach.

**Description of data:** This file contains the supplementary Table of Contents and Figures S1 through S11, illustrating the network topology, module evaluations, and dynamic rewiring across clades.

**File name:** Additional file 2

**File format:** .xlsx

**Title of data:** Supplementary Tables.

**Description of data:** This file contains Tables S1 through S5, providing detailed metadata for the 106 *E. coli* transcriptomic datasets, module functional annotations, and dataset-specific modularity performance metrics.

## Declarations

### Ethics approval and consent to participate

Not applicable

### Consent for publication

Not applicable.

### Availability of data and materials

The transcriptomic datasets underlying this article were obtained from public repositories and are available in the NCBI GEO database (https://www.ncbi.nlm.nih.gov/geo/). The exact dataset accession numbers (GSE identifiers) and corresponding metadata are detailed in Additional file 2, Table S1. The reference pan-genome data utilized in this study are available in the Microbial Genome Database (MBGD, https://mbgd.nibb.ac.jp/). Custom R script and code used for hypergraph-based pan-network construction, community detection, and downstream analysis are freely available at https://github.com/ginfolab/hyper_pan_network/

## Acknowledgements

Computational resources were provided by the Research Center for Computational Sciences (Project: NIBB, 24-IMS-C320, 25-IMS-C316, 26-IMS-C334).

## Authors’ contributions

I.U. conceptualized and supervised the project. Z.J. and I.U. designed the methodology. Z.J. developed the software, conducted the investigation, and performed the data visualization. Z.J. drafted the manuscript; I.U. reviewed and revised the manuscript. All authors read and approved the final manuscript.

