## Supplementary Figures for "Modular core network constructed from *Escherichia coli* transcriptome datasets using a hypergraph-based pan-network approach"

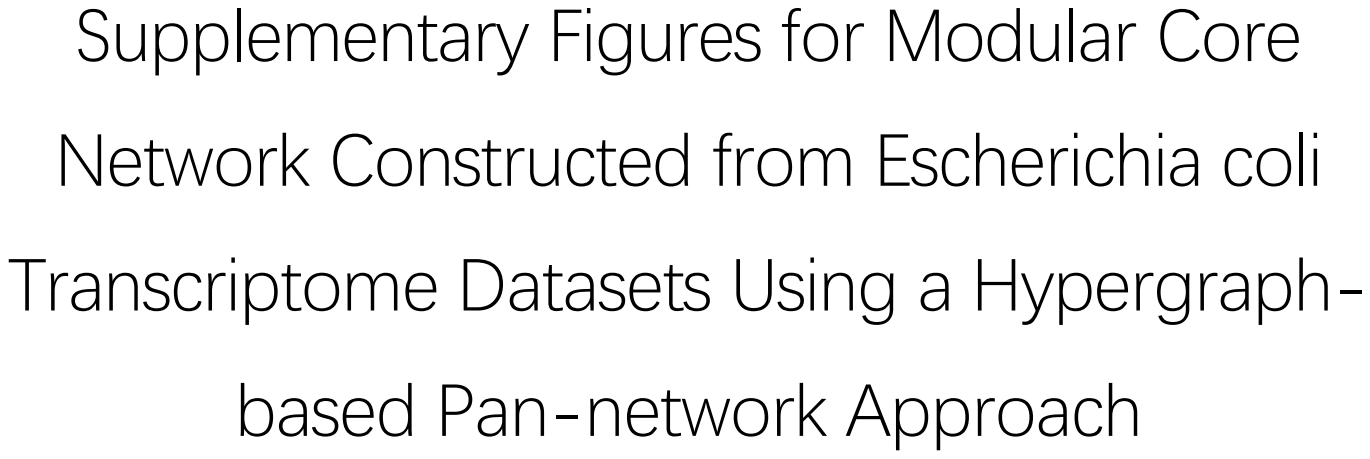

Supplementary Figures for Modular Core
Network Constructed from *Escherichia coli*
Transcriptome Datasets Using a Hypergraph-
based Pan-network Approach

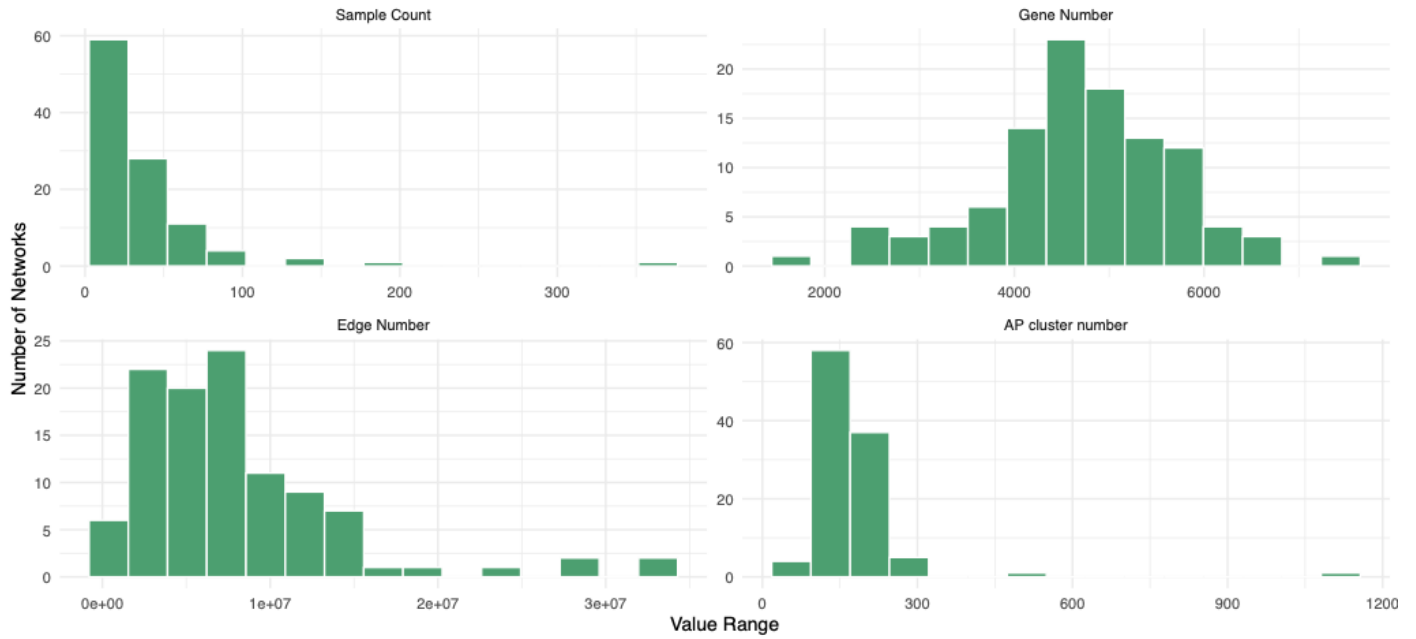

**Figure S1. Basic descriptive statistics of the 106 empirical co-expression networks.**

Basic statistics for 106 GSE series-based networks: Distributions of (A) the number of samples per dataset, (B) the number of expressed genes included in each network, (C) the total number of gene–gene edges derived from pairwise correlations, and (D) the number of clusters identified by AP clustering. These statistics illustrate the heterogeneity among datasets in sample size and network complexity.

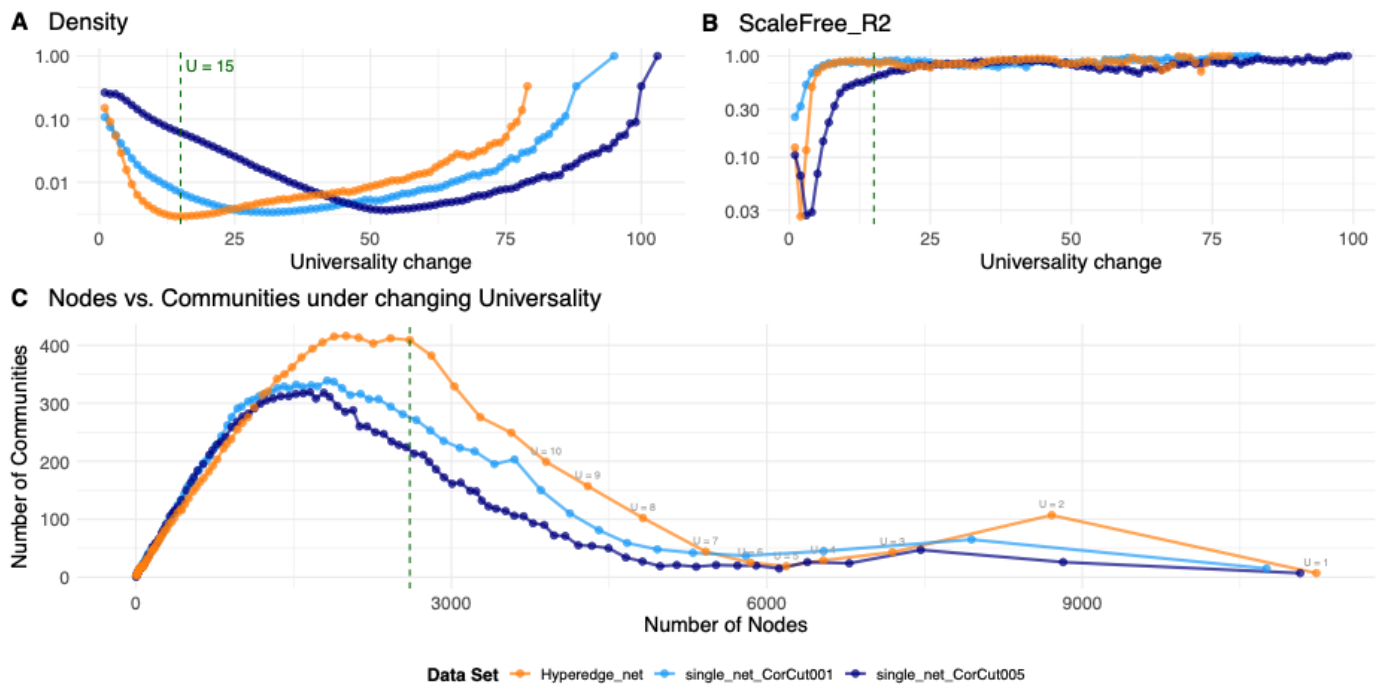

**Figure S2: Network Density, Scale-Free Fit, and Community Structure Under Changing Universality.**

Top panels: Network Density (left) and ScaleFree\_R<sup>2</sup> (right) for the three network types: Hyperedge\_net, single\_net\_CorCut001, and single\_net\_CorCut005, plotted across increasing U change. The green dashed line indicates U = 15, which marks the approximate point at which density and scale-free structure begin to stabilize. Bottom panel: Relationship between the number of nodes and the number of detected communities as U decreases from right to left. Labels show the corresponding U levels (U = 1, 2, 3, ...). The curves illustrate how network granularity and community fragmentation vary across different U constraints, with U = 15 serving as a reference stabilization point.

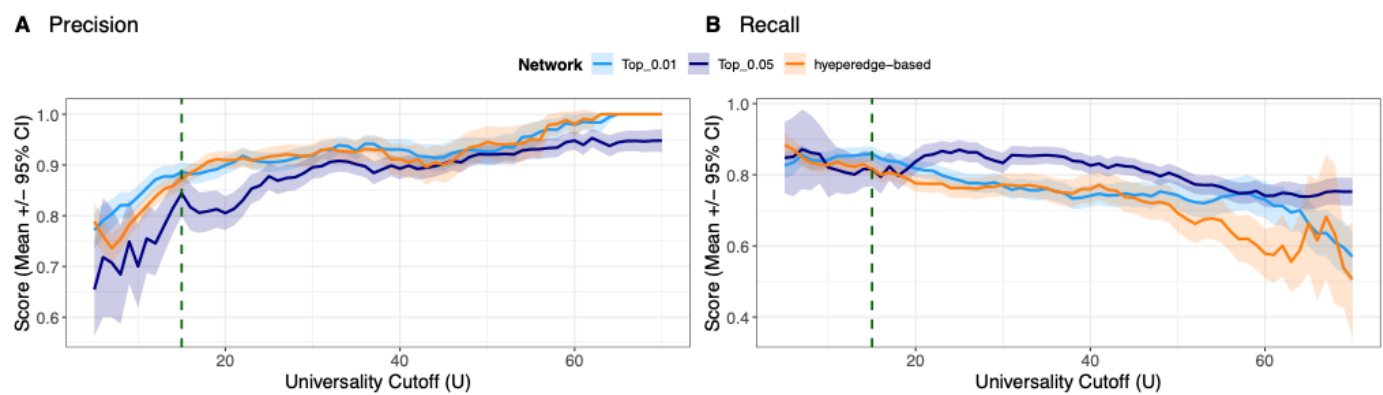

**Figure S3. Quality metrics for mapped operons across three network construction methods.**

Precision (A) and Recall (B) for mapped modules (F-score > 0.5) relative to RegulonDB plotted against increasing Universality (U) cutoffs for Hyperedge\_net, single\_net\_CorCut001, and single\_net\_CorCut005. Lines represent means with 95% confidence intervals. The green dashed line marks U = 15.

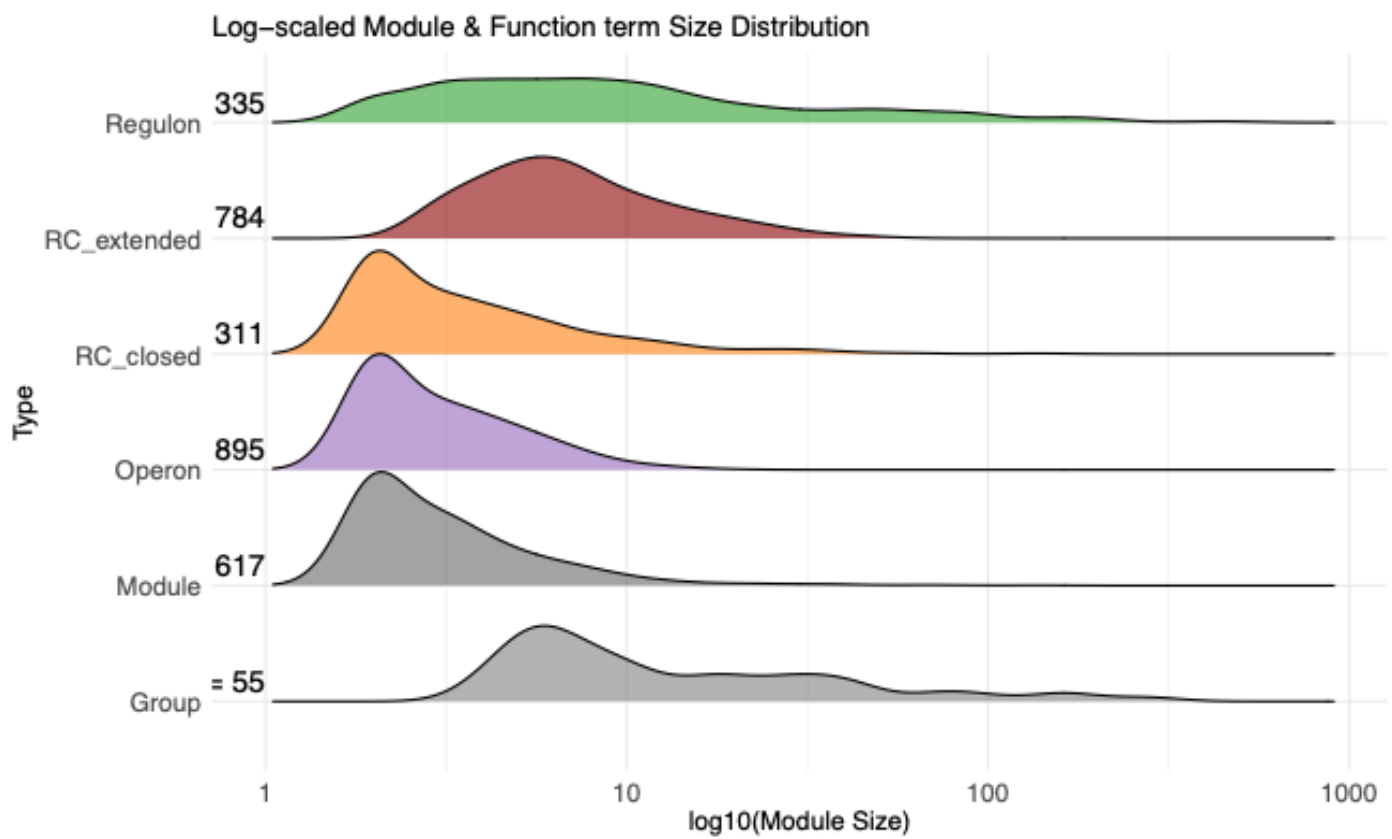

**Figure S4. Comparative size distribution of identified network communities and RegulonDB functional terms.**

The ridge plot displays the frequency distribution of gene counts (sizes) for the identified network communities (Module and Group) and standard regulatory units retrieved from RegulonDB (Regulon, Operon, etc.). The x-axis represents the term size on a log10 scale. The numbers listed to the left of each frequency curve indicate the total count of items in that specific category. The plot shows that the size distribution of Modules resembles that of Operons, whereas Groups exhibit a broader distribution, similar to Regulons.

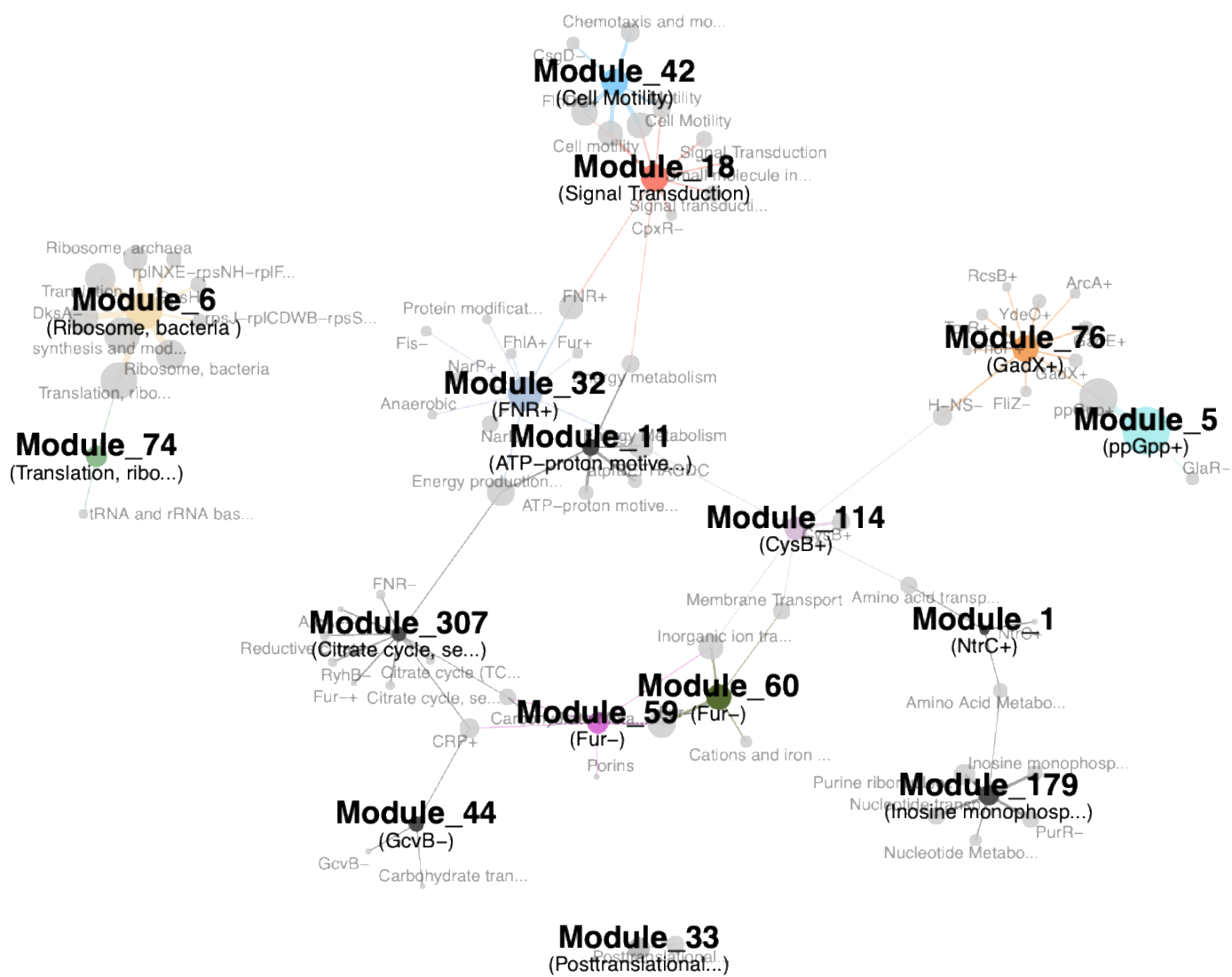

**Figure S5. Functional connectivity network of 16 representative modules.**

This bipartite network visualizes the topological relationships between modules and their enriched functions. The network comprises the top 11 modules ranked by gene count and 5 manually selected modules representing distinct functional groups. (1) Nodes: Large colored nodes represent the 16 selected modules, with size proportional to the total gene count within the module. Small grey nodes represent significant functional terms (derived from RegulonDB and COG, $q\text{-value} < 0.01$), sized by the number of genes involved. (2) Edges: Connections indicate significant enrichment. The edge thickness corresponds to the statistical significance ($-\log_{10}(q\text{-value})$). (3) Layout: The network uses a force-directed layout (Fruchterman-Reingold) where edge weights act as attractive forces, positioning functional terms spatially closer to their most strongly associated modules while maintaining distinct cluster separation.

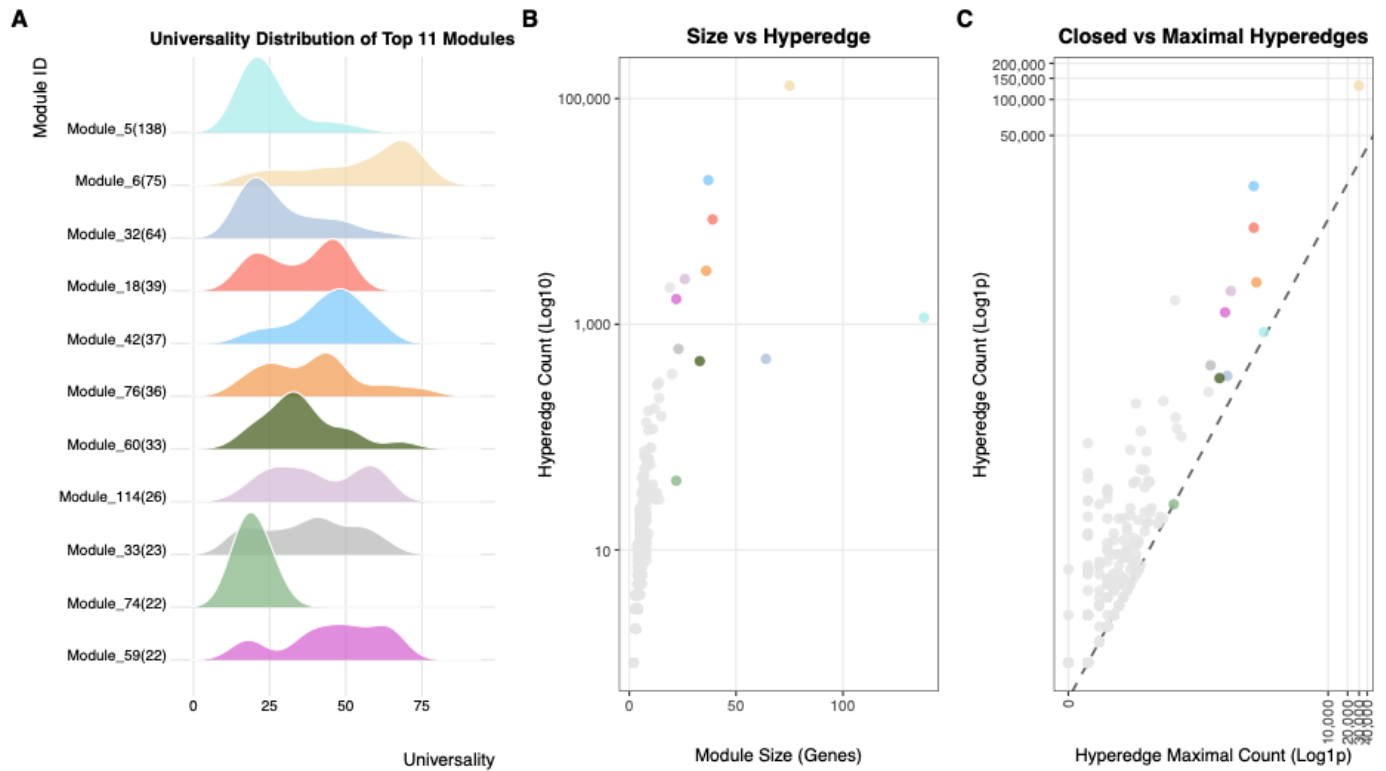

**Figure S6. Gene universality and structural properties of the core network modules.**

(A) Ridge plot displaying the frequency distribution of U values for genes across the top 11 largest modules identified in the core network. The y-axis lists the Module IDs (sorted in descending order by total gene count), with the exact number of genes included in parentheses. The x-axis represents the U score, quantifying the prevalence of each gene across datasets. (B) Scatter plot showing the relationship between module size (number of genes) and the corresponding number of closed hyperedges (log10 scale). (C) Scatter plot illustrating the relationship between the maximal itemset count (x-axis) and the hyperedge count (y-axis) for each module, both presented on a log1p scale. A dashed diagonal reference line ($y=x$) is included to indicate equality between the two counts. In panels (B) and (C), data points representing the top 11 modules are color-coded to match those in panel (A), while all other modules are shown in light gray.

### Top Regulators

### Structural Sub-Regions

### KEGG pathway

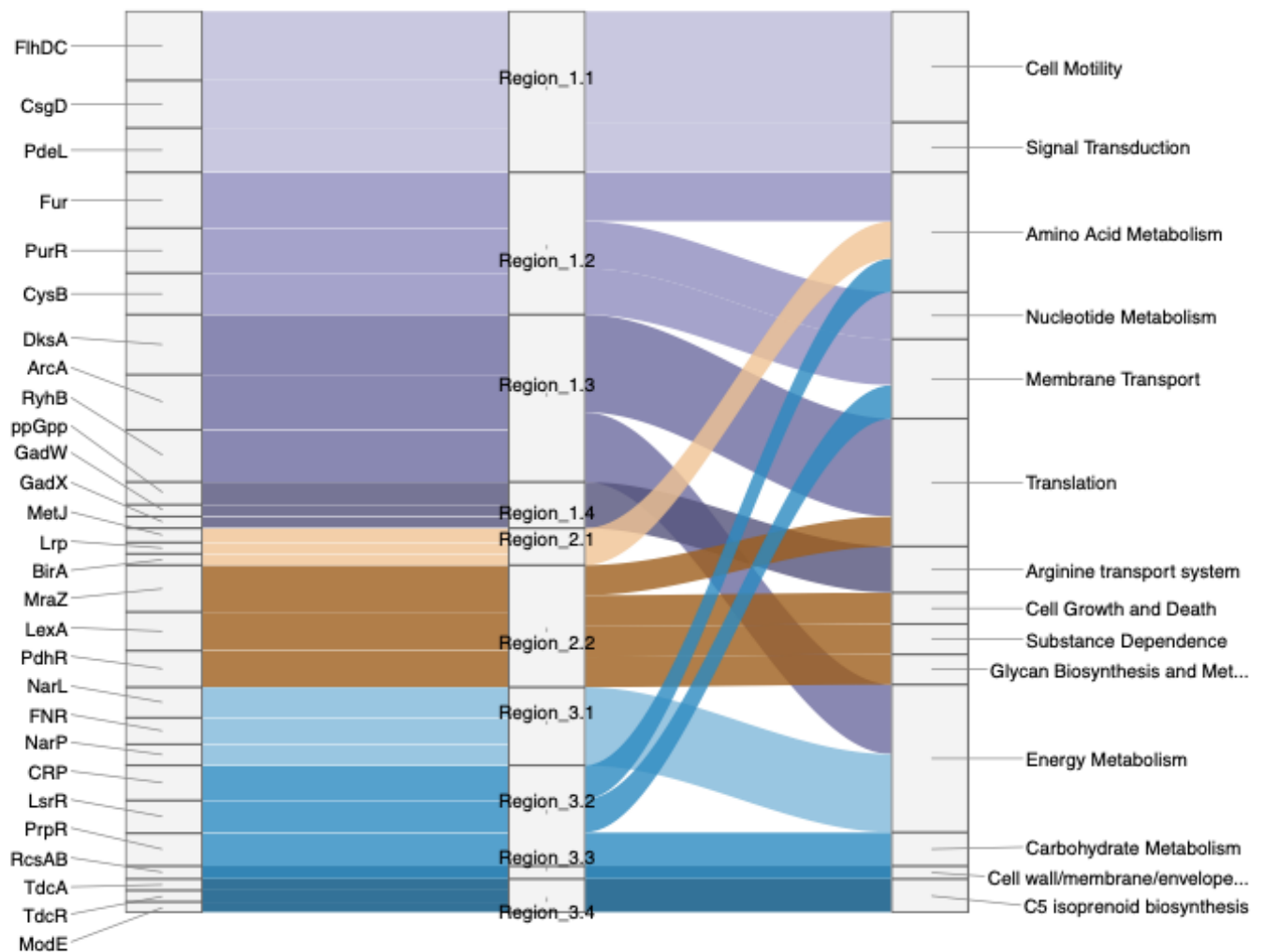

**Figure S7. Regulatory and functional information flow across the *E. coli* pan-network.**

The Sankey diagram illustrates the cascade from the top upstream regulators (left) to downstream phenotypic outputs (right, prioritized by KEGG annotations), with the latter bridged by structurally defined Subregions (middle). Connecting flow widths are proportional to the combined statistical significance ($-\log_{10}$ FDR) of regulatory interactions and functional enrichments. Note that the width is not proportional to the number of genes. Subregions are color-coded by parent domain (Region 1: purple; Region 2: orange; Region 3: blue), with intra-region gradient shading reflecting their hierarchical relationships.

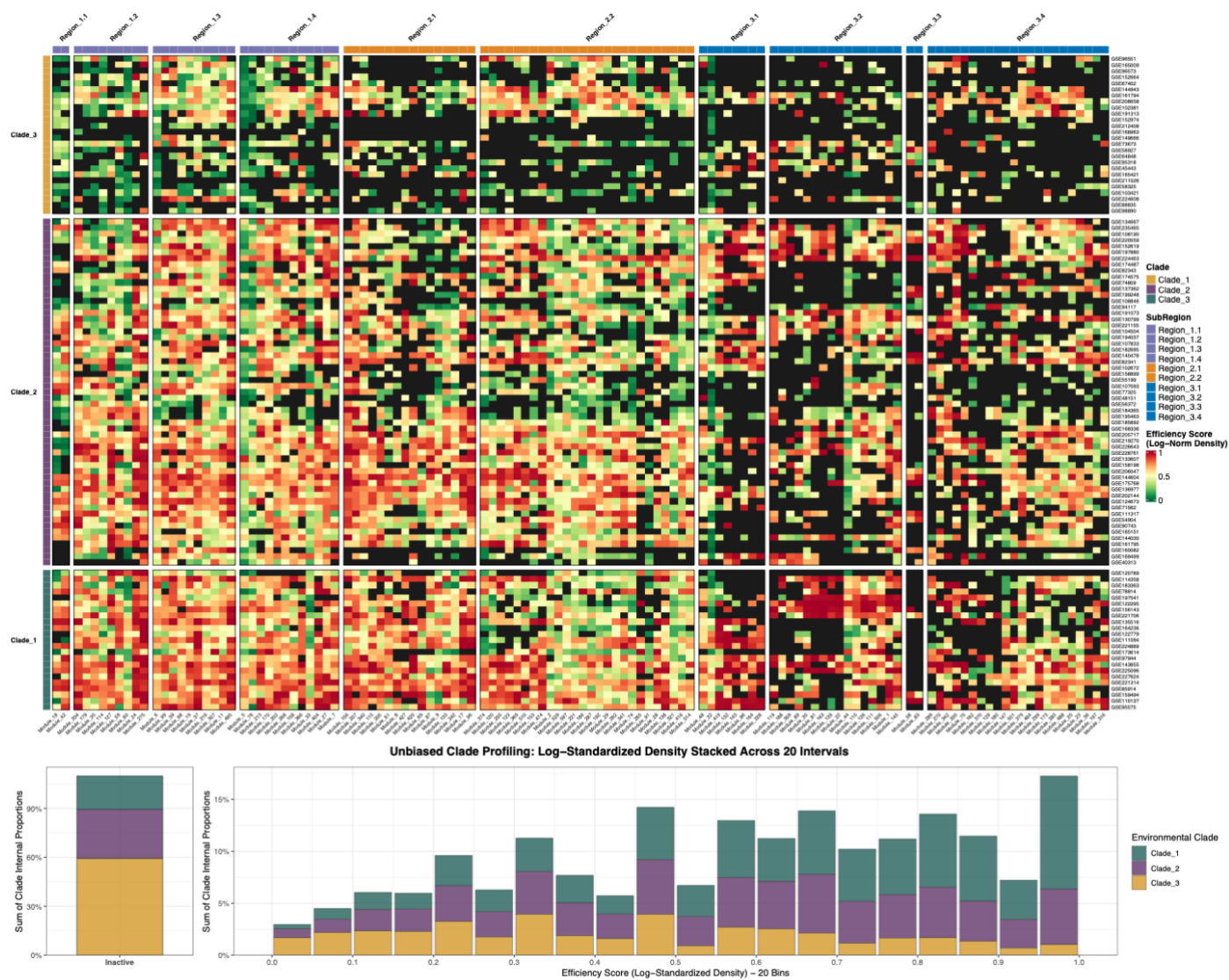

**Figure S8. The density profile of the *Escherichia coli* core-network modules.**

The heatmap displays the Log-Standardised Density of co-expression modules across datasets. Raw weighted densities were calculated based on the total internal interactions (combining **intra-module edges**) to comprehensively assess module tightness, then log-compressed to mitigate high-value polarization, and Min-Max normalized to a 0–1 scale. Color intensity transitions from green (low abundance) to red (highly dense topological cores). Rows and columns are structured by phylogenetic clades and subregions, respectively. The bottom panels present unbiased clade-specific density distributions. The y-axis represents internal proportions (module count normalized by total clade size) stacked across clades. The left bar shows the proportions of inactive modules, while the right histogram maps active modules across 20 density intervals, revealing shifts under varying environmental pressures.

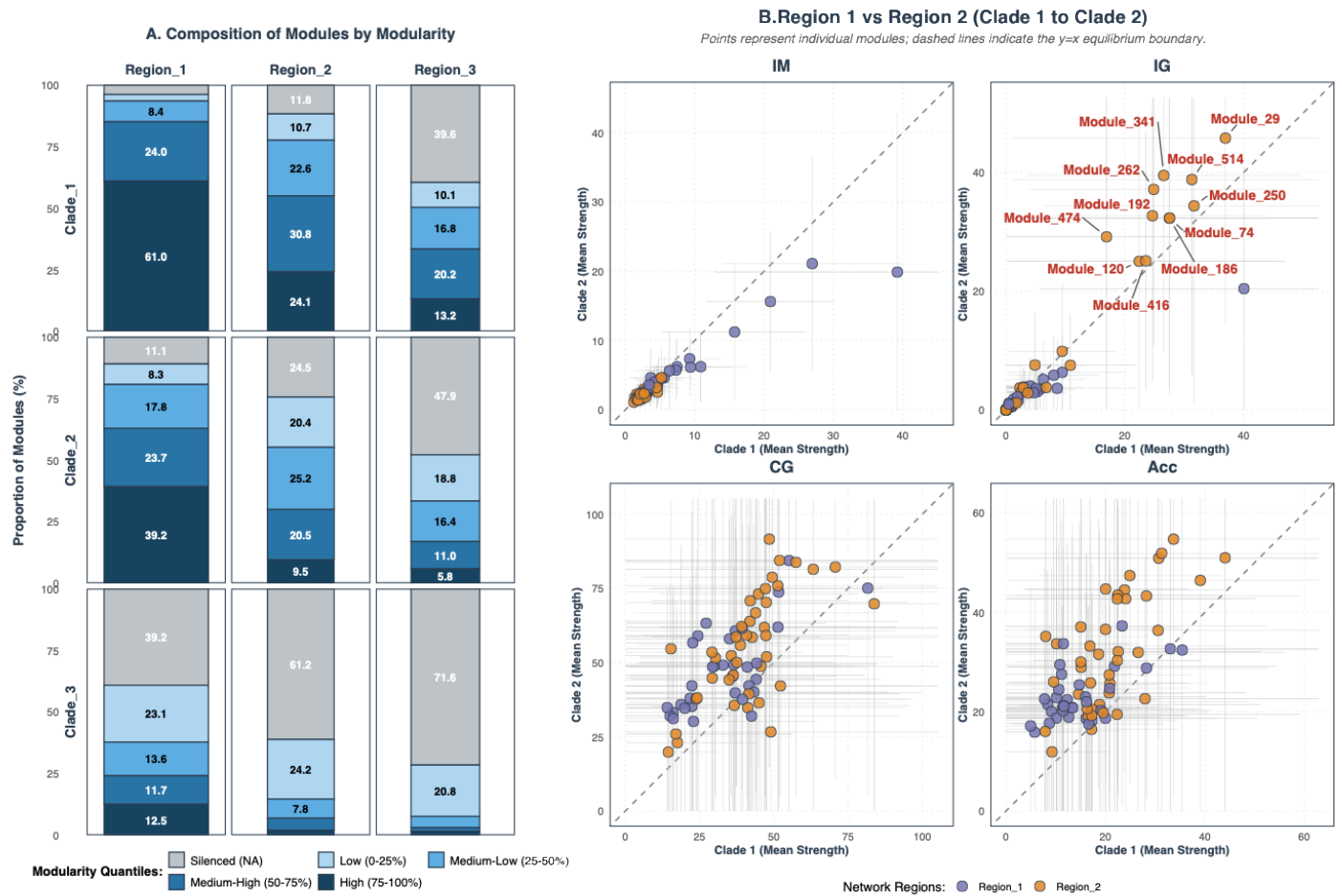

**Figure S9. Module independence and integration across environmental clades.**

(A) Module composition by quantiles reveals structural cohesiveness across environmental perturbations, stratified by macro-regions. Grey indicates inactivated/silenced states (valid nodes < 3 or modularity $Q_c \leq 0$), while the blue gradient shows proportions of activated modules grouped into modularity quartiles (Low [0-25%] to High [75-100%]). Bar width reflects the total gene count per macro-region. (B) Topological state transitions of activated modules from Clade 1 to Clade 2, segregated by four edge types (IM: Intra-module; IG: Intra-group; CG: Cross-group; Acc: Accessory). Each point represents a module's mean strength (the average number of connections per gene), mapped between Clade 1 (x-axis) and Clade 2 (y-axis). Colors denote macro-regions (Purple = Region 1; Orange = Region 2). The dashed diagonal ($y=x$) indicates whether connection strength increases (above) or decreases (below) under Clade 2 conditions. Error bars represent standard deviations across datasets within each clade, reflecting environmental plasticity. Labels in the IG subplot explicitly denote Group 14 modules.

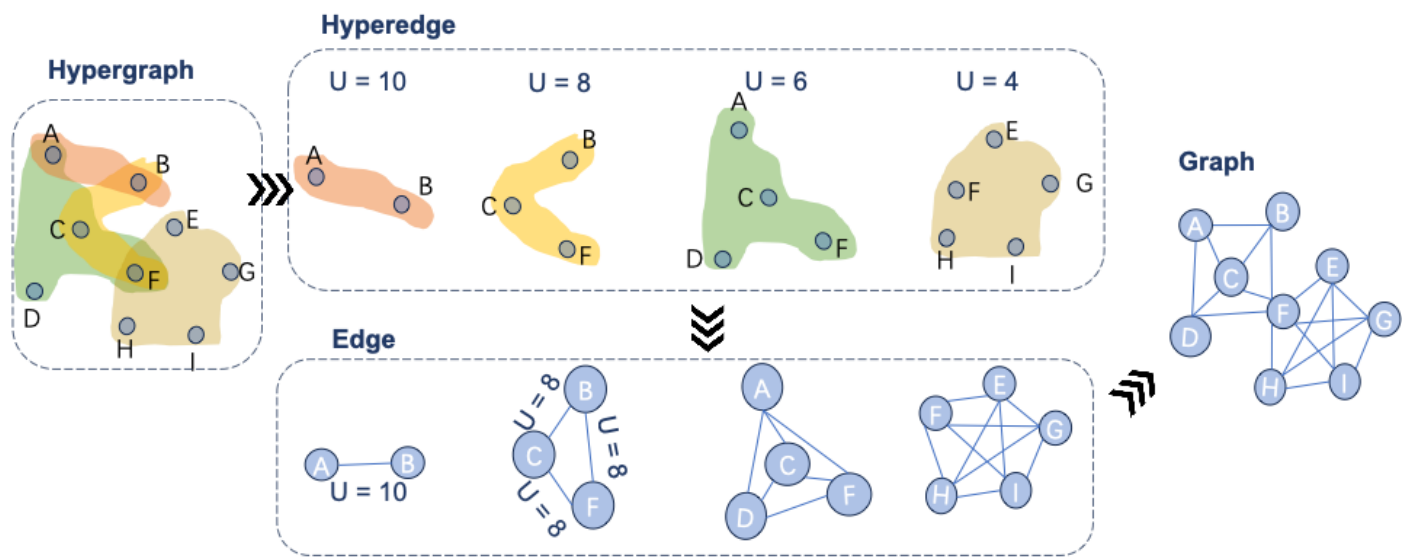

**Figure S10. Transformation of hyperedges into pairwise edges via clique expansion.**

Each hyperedge (colored region) represents a group of co-associated genes with a specific Universality ($U$). Hyperedges are decomposed into fully connected subgraphs (in the bottom panels). For gene pairs that occur in multiple hyperedges, the final assigned edge weight is the maximum $U$ value across the overlapping hyperedges.

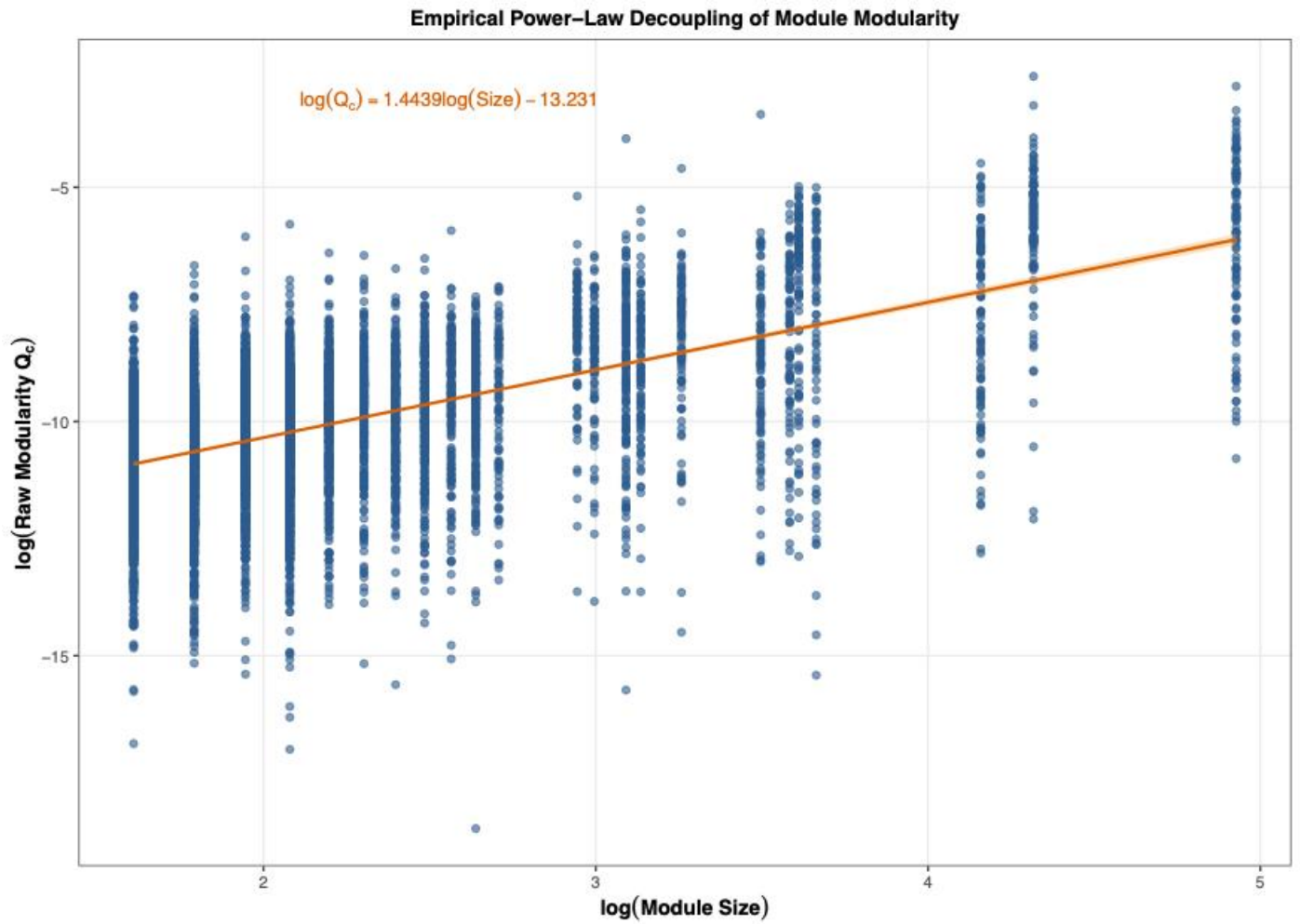

**Figure S11. Empirical power-law scaling between module size and raw modularity ($Q_c$).**

The log-log scatter plot illustrates the relationship between module size and raw modularity: $\log(Q_c)$ versus $\log(\text{Size})$.
